# Pond age and agricultural land cover influence the occurrence of *Anopheles maculipennis* in garden ponds in Hungary

**DOI:** 10.64898/2026.06.08.730838

**Authors:** Olivera Stamenković, Barbara Barta, Zsuzsanna Márton, Thu-Hương Huỳnh, Csilla Laskai, Andrew J. Hamer, Irene Tornero, László Zsolt Garamszegi, Márton Uhrin, Zsófia Horváth

## Abstract

Garden ponds are important elements of urban landscapes and provide numerous ecosystem services, but they may also support mosquito populations that can act as vectors of pathogens. However, mosquito occurrence in garden ponds has so far received limited attention. We engaged citizen scientists to provide data on their garden ponds and collect water samples for eDNA analysis to assess mosquito occurrence in 319 garden ponds across Hungary, with special focus on *Anopheles maculipennis,* a potential malaria-vector species. We tested the effects of land cover, local pond management activities and environmental factors on the probability of *An. maculipennis* occurrence in garden ponds. Overall mosquito occurrence was relatively low, with mosquitoes detected in 59 ponds (18%) and *An. maculipennis* detected in 46 ponds (14%). Agricultural land cover and pond age were associated with the occurrence of *An. maculipennis* in garden ponds. A higher percentage of agricultural land cover within 1000 m of garden ponds increased the probability of *An. maculipennis* occurrence, while occurrence was higher in newly created ponds (≤ 2 years). Our results suggest that landscape context is more important than local pond characteristics in determining the occurrence of *An. maculipennis* in garden ponds. This study contributes to a better understanding of the landscape context in which a potential malaria-vector species occurs in urban and peri-urban environments and highlights the value of citizen science for large-scale data collection from privately owned ponds that are otherwise inaccessible to researchers.

## Introduction

As the urban population is globally increasing, 68% of the human population is expected to reside in urban areas by 2050 (United Nations 2018), leading to further expansion of human-modified ecosystems, including urban and agricultural areas. Landscape anthropisation (i.e., urbanisation, deforestation, and agricultural development) leads to the loss of natural habitats and accelerates global warming, thereby filtering out sensitive taxa that cannot adapt to these environmental changes (Dirzo et al. 2014). In addition to biodiversity loss, other environmental issues can arise from human-driven landscape modification. For instance, altered environments can facilitate the emergence and spread of invasive species and certain mosquito vector species, which may generate health and economic problems in urban areas (Wilke et al. 2021).

While it is evident that landscape anthropisation has greatly contributed to the loss of natural habitats, including freshwater ecosystems (Burgin et al. 2016; Reid et al. 2019), garden ponds are increasing in numbers and are likely becoming the most frequent standing waterbodies in urban environments across Europe (Hill et al. 2021; Oertli et al. 2023). Garden ponds enhance aesthetic appeal, support biodiversity, and help regulate microclimatic conditions (Horváth et al. 2025). However, they may also promote higher mosquito abundance (Oertli and Parris 2019; Horváth et al. 2025), including species that may act as vectors of pathogens responsible for various diseases. The occurrence of specific mosquito species in aquatic habitats depends on various environmental factors, including physical and chemical water parameters, water depth, vegetation type, and hydrological conditions (Muturi et al. 2008; Yadav et al. 2012), as well as the presence of natural predators (Mokany and Shine 2003; Mataba et al. 2021) and habitat age (Munga et al. 2013). In addition, human-induced landscape alterations can affect mosquito distribution, usually favouring species of public health importance (Roiz et al. 2015; Perrin et al. 2022). To properly assess the risk of mosquito-borne diseases, it is essential to understand which landscape components affect mosquito occurrence and how their presence is related to local environmental factors.

Until the mid-twentieth century, malaria was a widespread disease in Europe. Over the last two decades, malaria reappeared in several countries in Southern Europe with high density of *Anopheles* species, driven by the increasing importation of infections through mass migration and travel (ECDC 2015, 2024). *Anopheles maculipennis* is considered the primary malaria vector in Europe (Bueno-Marí and Jiménez-Peydró 2011), and it is one of the most abundant (Tóth and Kenyeres 2012) and the most common mosquito species in Hungary (Soltész et al. 2025). While certain studies showed that this species is more common in non-urbanised areas (Pernat et al. 2021a; Duval et al. 2023), there is evidence that the occurrence of *Anopheles maculipennis* could be favoured by landscape anthropisation, in particular by agricultural land conversion (Ibáñez-Justicia et al. 2022; Gilioli et al. 2024). It may also prefer small, artificial or human-impacted habitats (Dakić et al. 2008; Kavran et al. 2018). However, there are limited data on its occurrence in garden ponds and factors regulating its distribution within these artificial habitats (Horváth et al. 2025). Besides local environmental factors (e.g. water chemistry, natural predators, vegetation), certain management activities practised by pond owners or specific garden pond features, such as water circulation, might also determine the probability of mosquito occurrence, and potentially repel mosquitoes, including *An. maculipennis* (EPA 2005; Rey et al. 2012; Horváth et al. 2025; Tornero et al. 2026). Still, the importance of these factors as predictors of mosquito occurrence in garden ponds remains poorly understood (Horváth et al. 2025). Identifying the factors that influence the use of garden ponds as breeding sites by malaria-vector *Anopheles* species is essential for developing effective control measures.

Citizen science has proven to be a useful tool for monitoring and collecting data on mosquito occurrence across broad spatial scales (Pernat et al. 2021b; Garamszegi et al. 2023). It can also be a useful approach for collecting data from garden ponds, as many of these artificial habitats are inaccessible to professional researchers since they are located on private properties (Horváth et al. 2025). Furthermore, the environmental DNA (eDNA) approach has been shown as an effective method for the detection of the most abundant mosquito species in stagnant waters from water samples (Schneider et al. 2016; Krol et al. 2019) since mosquito larvae often occur in high abundance, which increases the chance of DNA-based detection (Elbrecht et al. 2017; Krol et al. 2019). Additionally, juvenile stages of most mosquito species are associated with the water surface (Rejmánková et al. 2013), making their eDNA likely to be present in the upper part of the water column, where it is easily accessible for sampling (Krol et al. 2019). Finally, the eDNA approach enhances data accuracy and taxonomic resolution, both of which are often limited in citizen science observations (Horváth et al. 2025).

The objective of this study was to assess the probability of *An. maculipennis* occurrence in garden ponds across Hungary (Central Europe) in relation to landscape-level drivers, local environmental factors, and local management activities applied by pond owners and municipalities. Since central chemical mosquito-control treatment is regularly carried out in areas with the highest mosquito infestation, we also included the volume of chemical mosquito control (i.e., volume of insecticide spraying) as a potential driver of *An. maculipennis* occurrence in garden ponds. We hypothesise that landscape anthropisation, particularly agricultural land use, increases *An. maculipennis* occurrence in garden ponds, since higher occurrence of this species in agricultural areas has been reported previously (Ibáñez-Justicia et al. 2022; Gilioli et al. 2024). In contrast, we expect that the presence of predators (fish, odonates, amphibians) would reduce the occurrence of this species through predation on larvae. We also hypothesise that the probability of *An. maculipennis* occurrence would be lower in regularly managed garden ponds and in ponds with water-circulation elements. Overall, our findings can enhance the assessment of mosquito-borne disease risk at a national level.

## Methods

### Citizen science survey and study site selection

Garden ponds were selected from the database of the MyPond citizen science project (https://mypond.hu/en/), launched in 2021, which contains information on the biota, local features and management practices of garden ponds in Hungary (for details on the survey, see Hamer et al. 2024 and Márton et al. 2025). The data on pond biota used in this study included the pond owners’ observation of the frequency of occurrence of adult amphibians, tadpoles and odonates in and around ponds (0-never; 1-once; 2-sometimes; 3-often) and intentional introduction of fish, crayfish and turtles by pond owners (0 –not introduced; 1-introduced). Data on local features of ponds included pond age (years), length (cm), width (cm), maximum depth (m), substrate type (natural, concrete, rubber pond liners, or plastic), shoreline and surface vegetation cover (%) (see Hamer et al. 2024 for details). Pond age was transformed to a categorical variable: old ponds (>2 years old) and newly created (≤2 years old). These categories were selected because no precise data on pond age were available for all ponds, and previous research reported a significant difference in the biodiversity of garden ponds between these two age categories (Hamer et al. 2024; Márton et al. 2025). Finally, local management data of garden ponds included the use of insecticide/pesticide spray in the garden, pond draining, bottom cleaning, heating, the use of chemicals against mosquito larvae (pills) and water circulation (0-never, 1-less than once a year, 2-once a year, 3-more than once a year). Based on the pond’s width and length, we approximated the pond surface area (m^2^) by multiplying them. We acknowledge that this approach may introduce some overestimation for ponds with irregular shapes. We included only garden ponds with a surface area ≤ 200 m^2^ in the analysis, since most ponds larger than 200 m^2^ served as agricultural waterbodies (Hamer et al. 2024). We also excluded two ponds smaller than 200 m^2^ that were identified as agricultural farmland ponds. This resulted in a dataset of 319 garden ponds across the country (Fig. 1).

**Fig. 1.**
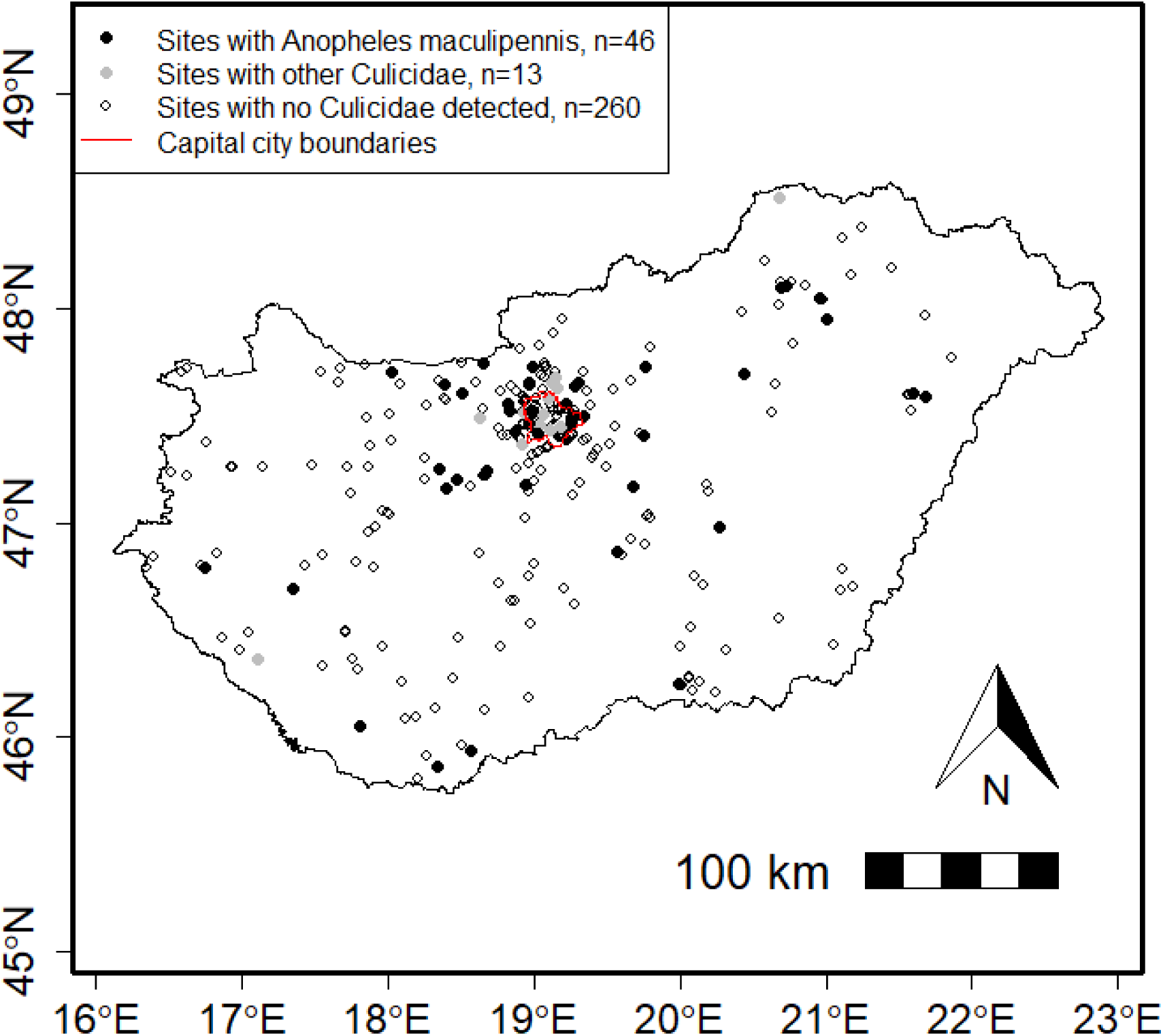
Distribution of sampling sites (garden ponds) across Hungary (n=319)

### Molecular methods

Water samples for eDNA analysis were collected from 319 garden ponds across Hungary in April and May 2022. Samples from 281 ponds were collected by pond owners, while samples from the remaining 38 ponds, located in Budapest, were collected by research teams following the same methodology as pond owners. Sampling kits with a handbook of instructions were provided to each pond owner, and a YouTube video explaining the eDNA sampling was created (https://www.youtube.com/watch?v=4anK98-WgQM&t=57s). Pond owners took water samples at arm’s reach (∼1 m) using a sterile ladle from the shoreline at a depth of 10 cm from ten different points to obtain a pooled sample. They pre-filtered the water using a 280 µm pore-sized homogenisation bag to filter out large plant parts or debris. They filtered the collected water samples using a 0.45 µm Sterivex filter and a sterile syringe. The instruction was to push as much water through the filter as possible, then dry the filter by pushing air through. Finally, they had to add 99% ethanol to preserve the samples during transport to the laboratory. The filtered samples were returned to our laboratory, where the ethanol was removed from the filters by pushing air through the filter cartridge using a sterile syringe. The samples were stored in a freezer until further processing at −20°C. DNA was extracted using a Qiagen DNeasy Blood and Tissue kit with an additional ZYMO PCR Inhibitor removal step. Extracted DNA samples (0.03 mL each) were shipped to EnviroDNA, Australia, for analysis.

Metabarcoding assay targeting cytochrome oxidase I (COI) gene was applied to detect Culicidae (ecul-coi) (Krol et al. 2019). Negative controls that consisted of extraction negatives, PCR negatives and positive controls were included during library construction. Library construction included two rounds of PCR, whereby the first round employed gene-specific primers to amplify the target region, and the second round incorporated sequencing adapters and unique barcodes for each sample-amplicon combination included in the library. Sequencing was carried out on Illumina sequencing platforms (iSeq 100 and NextSeq 2000). Following quality control filtering to remove primer sequences, truncated reads, and low-frequency reads, DNA sequences were clustered into Operational Taxonomic Units (OTUs) based on sequence similarity. Amplicon denoising, chimaera removal, and taxonomic assignment were performed with VSEARCH software (Rognes et al. 2016), whereby each amplicon sequence variant (ASV) was assigned a species identity using a threshold of 95% by comparing against a curated reference sequence database derived from the NCBI nt database. Where a species could not be assigned (i.e., reference database was deficient and/or taxa were poorly characterised), taxonomic assignments were vetted through BLASTN searches against NCBI (www.ncbi.nlm.nih.gov).

### Local environmental factors and chemical mosquito control

In addition to the data on ponds provided by pond owners, we measured water chemistry parameters of garden ponds in the laboratory. Water samples for water chemistry analyses were collected by pond owners or research teams in the same way as samples for eDNA analysis. After sampling, pond owners sent these water samples along with the eDNA samples. Chemistry parameters that were measured included: total organic carbon (mg/L; TOC) and total nitrogen (mg/L; TN) using a Multi N/C1903100 TC-TN analyser (Analytik Jena, Germany); total phosphorus (µg/L; TP), heavy metals (µg/L; Fe, Cu, Mn, Ni, Zn) and metalloid (µg/L; As) using the standard spectrophotometric methods following the protocols from Eaton et al. (2005), and pH and conductivity (µS/cm), measured by a HANNA multiprobe.

To quantify mosquito control intensity around each pond, we used municipality-level data on insecticide spraying conducted in Hungary between May and August 2021 (National Directorate General for Disaster Management 2023). The data from 2021 were used as a proxy as mosquito control in 2022 was applied after the eDNA sampling. Rather than representing a direct effect on mosquito occurrence in 2022, the 2021 data were used as a proxy for the relative intensity of mosquito control among municipalities. For this, we first joined the data on mosquito control volume on the municipality level to municipality polygons in QGIS v3.24.3 (QGIS Development Team 2022), resulting in a mosquito control volume value for each polygon. Then, we joined this resulting layer in QGIS with the garden pond coordinates to obtain the volume values for each pond.

### Land cover calculation

Land cover was assessed using the Ecosystem Map of Hungary (project KEHOP-430-VEKOP-15-2016-00001; Ministry of Agriculture 2019) in QGIS v3.24.3 (QGIS Development Team 2022). First, we counted the number of 20×20 m raster pixels that correspond to six different land cover types: urban land cover (including short and tall buildings, transportation infrastructure and other artificial surfaces), green space in urban area, agricultural land, grasslands, forests, and permanent waters (including wetlands, lakes and other standing waters, and rivers and other running waters) within a 1-km buffer zone around each pond, by using Zonal Histogram tool. Then we calculated the proportion of total pixels within the buffers for each land cover category. In the same way, we calculated temporal water cover as an additional land cover category that includes smaller patches of temporary water, using the Copernicus High Resolution Water and Wetness (WAW) 2018 layer (CLMS 2020) in QGIS, as these are unavailable on the Ecosystem Map of Hungary.

### Data analysis

To analyse which land cover category, local environmental parameters and management types had a significant effect on the occurrence of *An. maculipennis* in garden ponds, we used generalised linear models (GLM) fitted with a binomial distribution and logit link function using the ‘glm2’ package in R (Marschner 2018).

In the first step, we performed a principal component analysis (PCA) on local environmental parameters (i.e., water chemistry), including pH, conductivity, TN, TP, heavy metals and As, using the *prcomp* function in the ‘vegan’ package in R (Oksanen et al., 2022), to reduce the number of predictors in the models. We extracted the site scores of the first two principal components, which explained 33% of the variation in water chemistry: PC1 (high negative loadings for heavy metals: Ni, Fe, Mn), and PC2 (high negative loadings for pH and conductivity), and used them as separate variables in further analysis. Additionally, we excluded predictors with low variability, including crayfish and turtle presence, heating and the use of chemicals against mosquito larvae (see Figs. S1, S2 and S3). Although the use of pills against mosquito larvae could have effects on mosquito occurrence patterns, we did not have much data on their use, with only one positive report of their application in the studied garden ponds (Fig. S2)

To assess possible associations among variables, we conducted Multiple Factor Analysis (MFA) using the packages ‘FactoMineR’ (Husson et al. 2023) and ‘factoextra’ (Kassambara and Mundt 2020). This allowed us to combine continuous variables with binary and ordinal variables with continuous ones in an unconstrained ordination. We visualised general patterns and associations between mosquito presence and 21 quantitative variables grouped into seven categories: land cover, temporary water bodies, pond management, chemical mosquito control, predators, water chemistry, and pond morphometry (Fig. S4). Based on MFA (Fig. S4), we selected ten variables that could potentially predict the probability of *An. maculipennis* presence, to be included as predictors in the model, and divided them into five groups: land cover (urbanisation, green space in urban area, agriculture), local pond management activities (draining, bottom cleaning, leaves removal and the use of insecticide/pesticide spray in the garden), chemical mosquito control volume, natural predators (tadpoles), and water chemistry parameters (PC2). Additionally, we performed pairwise Spearman’s rank correlation of the predictor variables (Fig. S5), to avoid using correlated variables in our models. Grasslands and forests were not included in the model as they were highly correlated with urbanisation (*ρ* > 0.7; Fig. S5). In addition to the selected quantitative variables, we included qualitative variables as predictors: pond age, vegetation cover (surface and shoreline) and bottom material. To test for potential non-linear relationships between *An. maculipennis* presence and land cover categories, we added quadratic terms to the corresponding predictor variables.

After selecting predictor variables, we performed model selection based on Akaike’s Information Criteria (AIC) using the ‘performance’ package in R (Lüdecke et al. 2025), including a null model (with no predictors), and considering all models with a ΔAIC ≤ 2 to have substantial support (Burnham and Anderson 2002). Initially, we developed eight models based on a combination of different predictor groups. Our full model included all groups: land cover + pond management + mosquito control + predators + water chemistry + pond age + vegetation + bottom material. The remaining seven models were created by excluding one group from each subsequent model (see details in Table S1). The simplest model only included land cover categories as predictor variables (Table S1). Before model selection, full models were tested for multicollinearity by applying variance inflation factors (VIFs) using the *vif* function from the R package ‘car’ (Fox and Weisberg 2019). After model selection, non-significant quadratic terms were excluded from the best model (model with the lowest AIC), and the new model was kept if excluding quadratic terms improved model fit (indicated by lower AIC). Finally, we checked for spatial autocorrelation of the residuals of the final selected model by using the ‘DHARMa’ package (Hartig 2024). First, we generated scaled quantile residuals via 1000 simulations from the fitted GLM. Then, we tested for spatial autocorrelation using the *testSpatialAutocorrelation* function, which calculates a Global Moran’s I statistic based on a Euclidean distance matrix of the observation coordinates. Statistical significance was determined by comparing the observed Moran’s I to the expected value under the null hypothesis of spatial randomness.

Visualisation of GLM results (effect sizes) was carried out using the ‘jtools’ package in R (Long 2024).

In addition, we tested the differences of selected environmental variables (conductivity, TOC, TP, TN, pond depth, tadpoles, adult amphibians, odonates, fish) between the two pond age categories in order to explain the obtained results of GLM. Since preliminary testing confirmed violation of normality assumptions, we performed non-parametric Kruskal-Wallis rank sum tests, followed by post-hoc Dunn’s tests with a Bonferroni correction, using the ‘FSA’ R package (Ogle et al. 2026). Results were visualised by using ‘ggplot2’ (Wickham 2016).

All statistical analyses were performed in R version 4.3.1 (R Core Team 2023).

## Results

Species from the Culicidae family were recorded in 59 out of 319 selected garden ponds, of which *An. maculipennis* was recorded in 46 garden ponds (Fig. 1). The remaining mosquito species were: *Aedes vexans* (recorded in three garden ponds), *Culex hortensis* (recorded in eight garden ponds), *Culex* sp. (recorded in one garden pond) and *Culiseta* sp. (recorded in six garden ponds). The remaining detected mosquito species were present only in a small number of the studied garden ponds; therefore, their occurrence patterns were not analysed separately.

The model with the lowest AIC (AIC = 257.5) included land cover categories (urbanisation, green space in urban area, agriculture), pond management (draining, bottom cleaning, leaves removal and the use of insecticide/pesticide spray), mosquito control volume, predators (tadpoles), water chemistry (PC2) and pond age (Table S1). After excluding non-significant quadratic terms from the model, the model fit was improved (AIC = 252.6; Table S1). Therefore, the model without quadratic terms of the land cover categories was selected as the final one. No spatial autocorrelation of the residuals was detected (observed = 0.0019; expected = −0.0033; p = 0.7887), suggesting that the model adequately accounted for the spatial structure of the data.

Agricultural land cover was a significant predictor of *An. maculipennis* occurrence (Fig. 2; Table S2), showing that there was a higher probability of *An. maculipennis* occurrence with increasing levels of agricultural land cover (Figs. 2 and 3). Pond age was another significant predictor of *An. maculipennis* occurrence (Fig. 2; Table S2) with a higher probability of its occurrence in newly created ponds (≤ 2 years) (Fig. 3). We did not detect any significant effects of local pond management activities, mosquito control, water chemistry or predators on *An. maculipennis* occurrence (Fig. 2; Table S2).

**Fig. 2.**
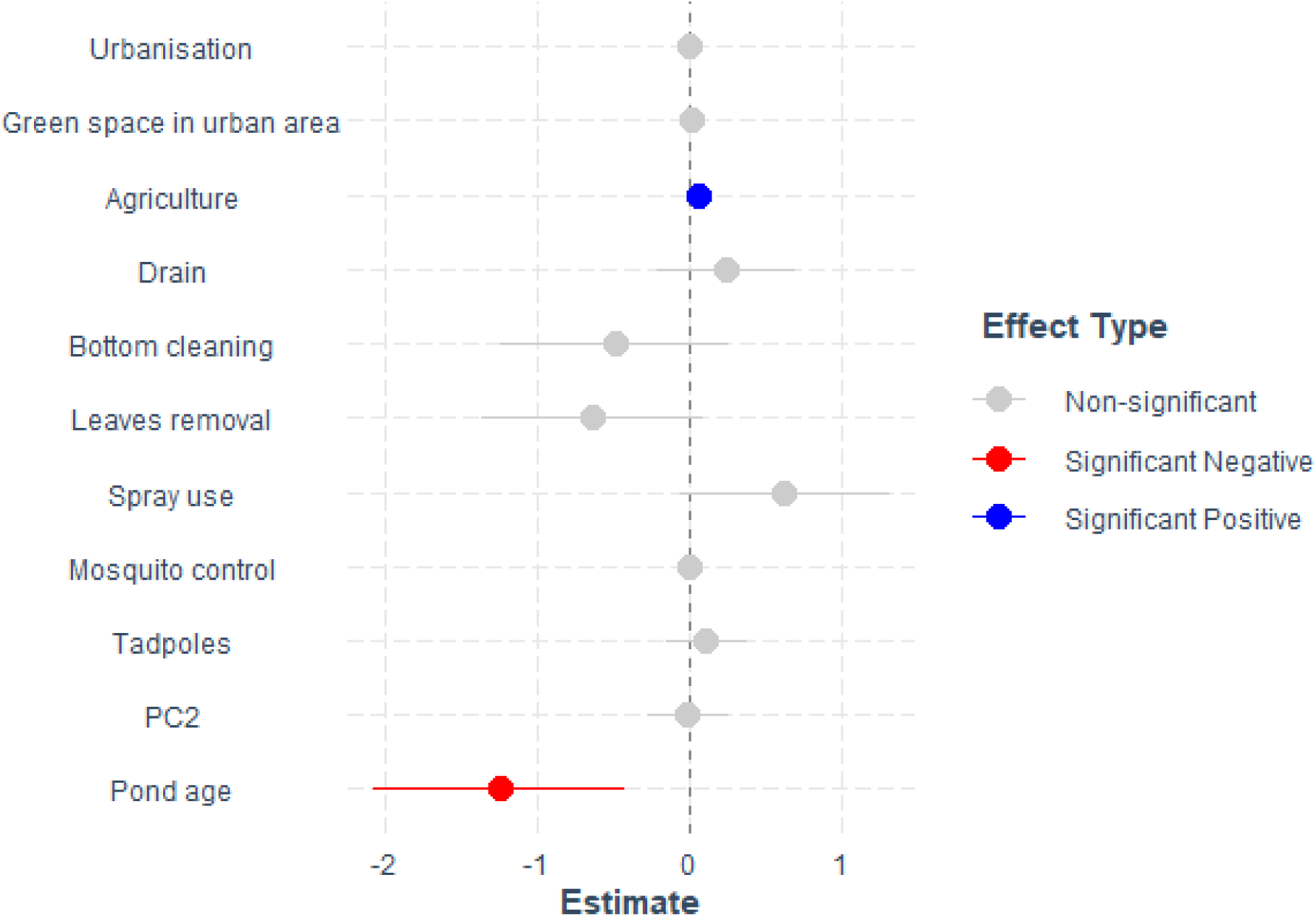
Effect size (coefficient estimates ± SE from the final selected generalised linear model) of each predictor on the probability of the occurrence of *Anopheles maculipennis* in garden ponds. Significant effects (p-value < 0.05) are indicated by different colours. PC2: second principal component (high negative loadings for pH and conductivity)

**Fig. 3.**
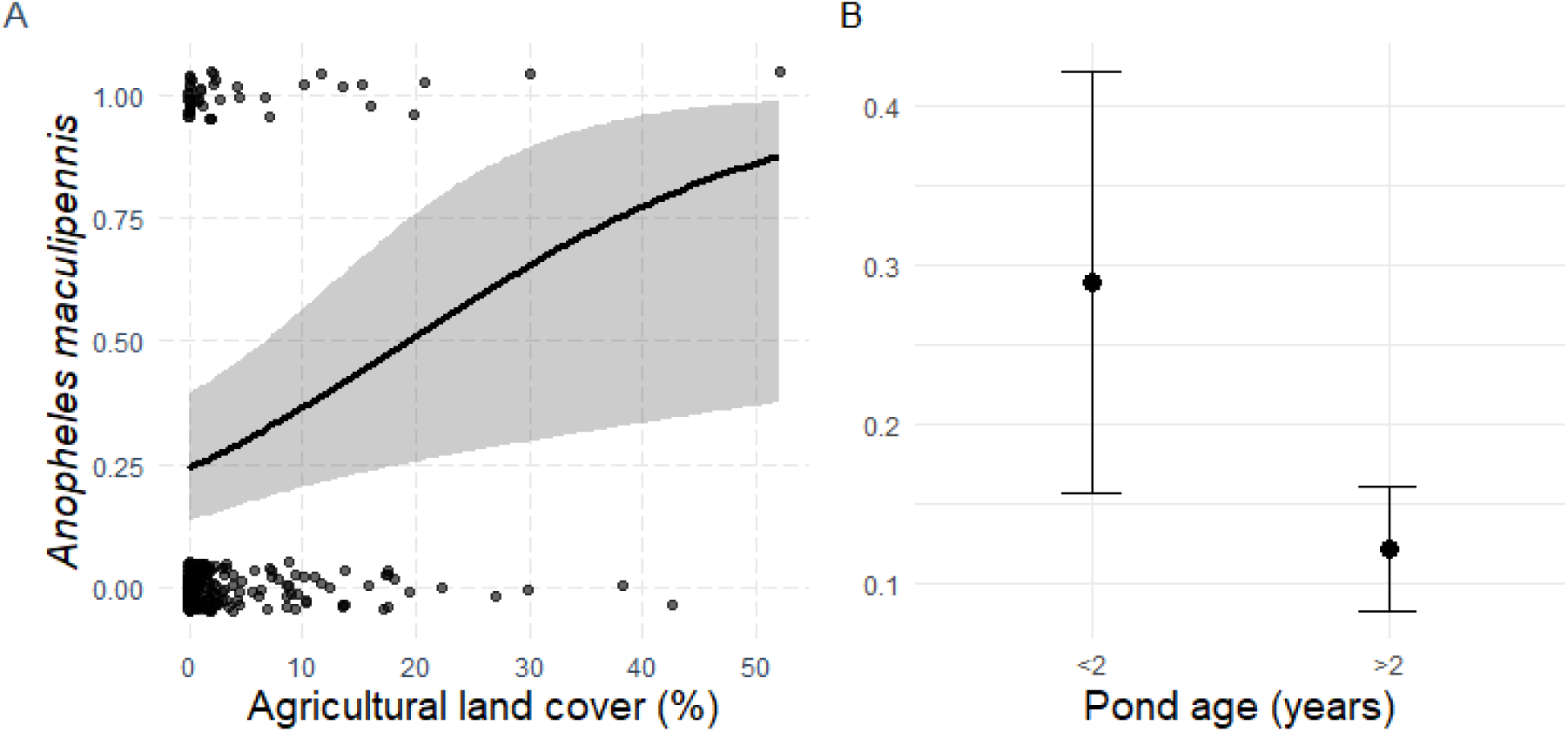
Effects of agricultural land cover (A) and pond age (B) on the probability of the occurrence of *Anopheles maculipennis* in garden ponds. For agricultural land cover (A), the curve represents predicted values from the final generalised linear model with binomial distribution. Study ponds are displayed with black dots. Gray shading represents the 95% confidence interval. For pond age (B), interval plots show predicted values of *An. maculipennis* occurrence probability in garden ponds of two different age categories: newly created ponds (≤ 2 years old) and old ponds (> 2 years old), derived from the same generalised linear model

## Discussion

Increased mosquito populations have been recognised as a typical ecosystem disservice of garden ponds (Oertli and Parris 2019; Horváth et al. 2025). However, only a limited number of studies examined the occurrence of mosquitoes in garden ponds and the factors that might influence their use of these habitats (Mokany and Shine 2003; Perea and Callaghan 2017). To our knowledge, this is the first study that utilised citizen-collected eDNA data to detect mosquitoes from garden ponds on a countrywide level. By examining the effects of land use, local management activities and environmental factors on the *Anopheles maculipennis* occurrence probability in garden ponds, our study improves our understanding of the habitat use of this potential malaria vector species in garden ponds. Focusing on *Anopheles maculipennis*, a potential malaria vector, we found that occurrence probability was primarily decreasing with pond age, and increasing with the proportion of surrounding agricultural land, while most local environmental and management variables had little explanatory power.

Pond age was the most important pond feature determining the occurrence of *An. maculipennis* in garden ponds, which is consistent with previous findings highlighting pond age as an important predictor of animal taxa occurrence in garden ponds (Hamer et al. 2024; Márton et al. 2025). The probability of the occurrence of *An. maculipennis* in garden ponds was significantly lower in ponds older than two years. Higher pond age may reflect later stages of successional processes in garden ponds, including complex changes to habitat structure, community composition and biotic interactions. Related to this, as ponds mature, they generally support higher biodiversity and increasingly complex communities, including a wider range of potential mosquito predators (Mari et al. 2010). In particular, garden ponds older than two years are more likely to contain fish introduced by pond owners, as reported by Hamer et al. (2024), who analysed a larger dataset of more than 700 garden ponds collected through the same citizen science program. Similarly, Márton et al. (2025) reported that newly created garden ponds (≤ 2 years old) were less likely to be inhabited by adult amphibians, tadpoles and odonates. The patterns observed in our eDNA subset were consistent with those reported by Márton et al. (2025), with tadpoles and adult amphibians occurring more frequently in older ponds (Fig. S6). However, neither tadpoles nor adult amphibians were significant predictors of *An. maculipennis* occurrence in our models. Furthermore, we found no significant differences in key abiotic characteristics (TOC, conductivity, TP, TN and pond depth) between the two pond-age categories (Fig. S6). These results suggest that the relationship between pond age and *An. maculipennis* occurrence is unlikely to be explained by a single environmental factor. Instead, pond age may act as an integrative proxy for ecological succession, reflecting changes in community composition, species interactions and habitat complexity that were not fully captured by the variables measured in this study. Future studies should also consider the effects of other mosquito predators, such as aquatic Coleoptera and Heteroptera, on the occurrence of *Anopheles* and other mosquito larvae, as these insects can also have an important role in mosquito control (Onen et al. 2024), and these groups increase in diversity with pond ageing (e.g., Mari et al. 2010).

Our results showed that agricultural land cover is an important factor in predicting the occurrence of *An. maculipennis* in garden ponds. The positive association of *Anopheles* species occurrence with agricultural land cover has been recorded previously by numerous studies (Kavran et al. 2018; Zeru et al. 2020; Mataba et al. 2021; Mburu et al. 2021; Perrin et al. 2023). This has usually been attributed to the increased availability of potential breeding habitats for mosquitoes (irrigation facilities), but also to the presence of cattle that provide sufficient blood meal and thus enhance mosquito reproductive success, or to the presence of animal shelters, which could be used as resting sites (Kavran et al. 2018; Zeru et al. 2020; Mburu et al. 2021). This pattern indicates that garden ponds may be used by *An. maculipennis* primarily in landscapes where source populations are supported by surrounding agricultural habitats, which may provide alternative breeding habitats, resting sites, and suitable vertebrate hosts in terms of domestic animals (Danabalan et al. 2014; Kavran et al. 2018).

Our study did not provide any evidence for the effects of local management activities practised by pond owners on the occurrence of *An. maculipennis* in garden ponds. Although insecticides can negatively affect aquatic insects (Ito et al. 2020), we detected no association between the use of insecticide or pesticide sprays and *An. maculipennis* occurrence. This may reflect the limited transfer of garden-applied chemicals into ponds, which are often isolated from polluted surface runoff (Márton et al. 2025). Similarly, we found no significant effects of bottom cleaning, draining or leaf litter removal from garden ponds, even though these activities are generally recommended in case of small artificial habitats to minimise mosquito breeding (EPA 2005). In case of draining, the frequency and timing of draining events may be more important than whether a pond is drained at all, given that mosquito larvae require a water habitat that exists for a minimum of seven days (EPA 2005; Henn et al. 2008), which is substantially shorter than the typical frequency of pond draining reported by garden pond owners. Consequently, even regularly drained ponds may provide suitable habitat for *An. maculipennis*. Similar to local pond management activities, we did not detect a significant effect of mosquito control on *An. maculipennis* occurrence. Previous studies have reported that chemical mosquito control can be effective in reducing mosquito densities for a short period of time (Garamszegi et al. 2026), but it is not effective in suppressing the entire mosquito population over the long term (Manica et al. 2017). As municipality-level mosquito control data from 2021 were used as a proxy for relative differences in mosquito control effort rather than direct exposure during the sampling year, the absence of a detectable effect should be interpreted cautiously. Nevertheless, our results suggest that mosquito control intensity was a poor predictor of *An. maculipennis* occurrence in garden ponds. Although we detected no association between local management activities and *An. maculipennis* occurrence, these practices may still influence mosquito abundance within ponds, which we could not assess here, as our metabarcoding approach was designed to assess occurrence rather than abundance.

## Conclusion

Our study provides the first countrywide assessment of mosquito occurrence in garden ponds using citizen-collected eDNA data, with a particular focus on *Anopheles maculipennis*, a potential malaria-vector species. Overall, mosquito occurrence in garden ponds was relatively low (in 59 ponds, 18%), with *An. maculipennis* being detected in only 46 of 319 ponds (14%). No invasive mosquito species were detected, suggesting that garden ponds are unlikely to play a major role in the current spread of invasive mosquitoes in Hungary. Our results revealed that the surrounding landscape of a garden pond and its age significantly affect the probability of *An. maculipennis* occurrence, with a higher probability in newly created garden ponds, and in those surrounded by agricultural land. Furthermore, our study highlights the value of citizen science approach in garden pond research, enabling large-scale data collection that would have otherwise been difficult to achieve. Future studies incorporating repeated seasonal sampling could provide more insightful results as many mosquito species achieve peak in their abundances in late spring or summer (Montarsi et al. 2015).

## Supporting information

Suppementary material

## Acknowledgements

We are thankful to the pond owners who provided the data on their garden ponds and collected the samples for eDNA analysis. We acknowledge those who helped in the field and laboratory work, website construction and correspondence with citizens: Beáta Szabó, Bence Gergácz, Bence Buttyán, Bernadette Szabó, Kata Bene, Vivien Kardos, Lilla Kocsis, and Dorina Varga. This research was conducted under permit FPH059/1107-2-2022 authorised by the Municipality of Budapest.

## Authors’ contributions

**Olivera Stamenković:** conceptualisation, formal analysis, visualisation, writing - original draft. **Barbara Barta:** investigation, project administration, visualisation, data curation, writing - review and editing. **Zsuzsanna Márton:**investigation, project administration, writing - review and editing. **Thu-Hương Huỳnh:** investigation, writing - review and editing. **Csilla Laskai:** investigation, project administration, writing - review and editing. **Andrew Hamer:** investigation, writing - review and editing. **Irene Tornero:** investigation, writing - review and editing. **László Zsolt Garamszegi:** resources, writing - review and editing. **Márton Uhrin:** writing - review and editing. **Zsófia Horváth:** conceptualisation, resources, investigation, funding acquisition, project administration, writing - review and editing.

## Availability of data and materials

All the data generated and analysed during this study are available upon request.

