## Supplementary material for "Pond age and agricultural land cover influence the occurrence of *Anopheles maculipennis* in garden ponds in Hungary": Suppementary material

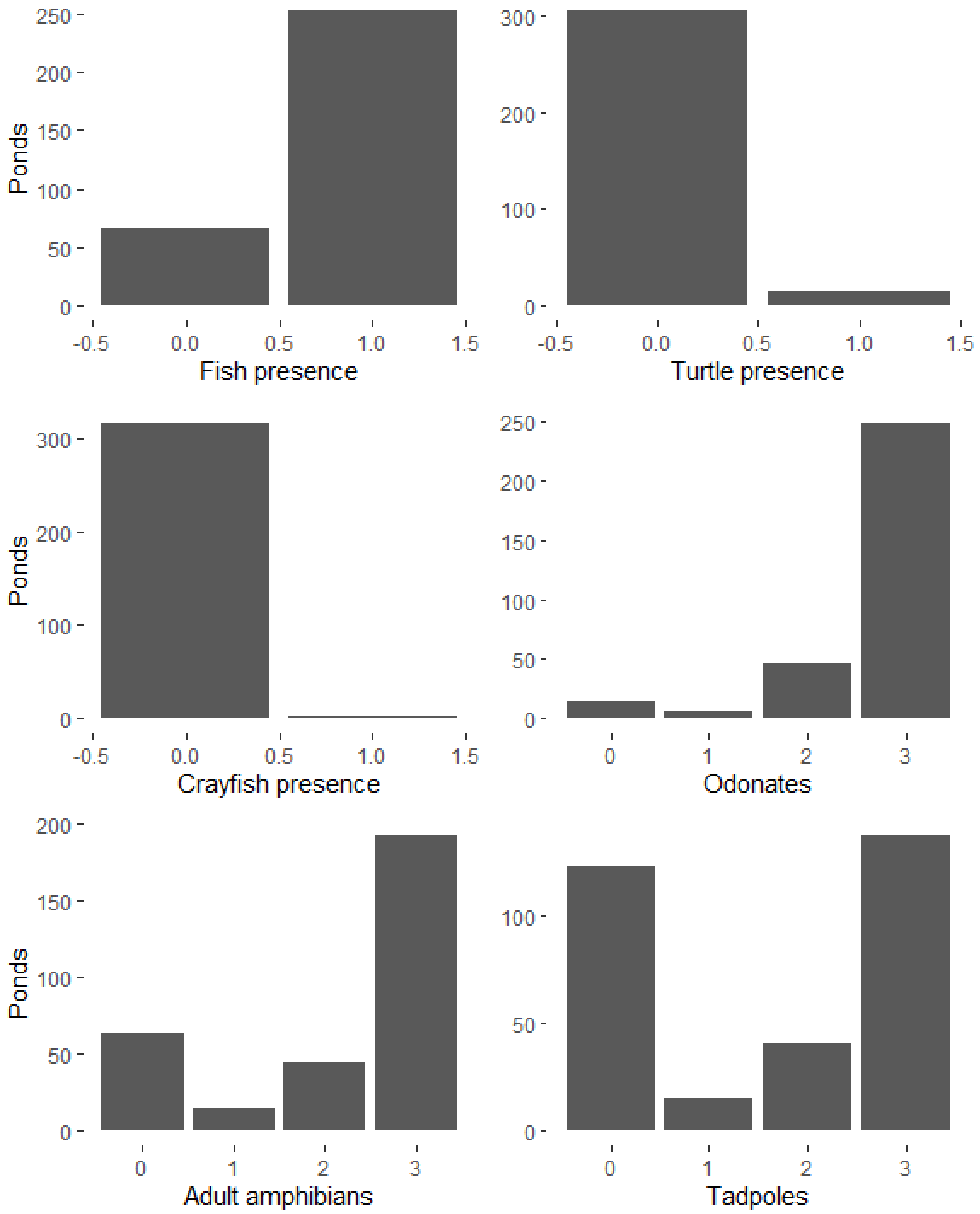


**Fig. S1** Bar plots showing the count of garden ponds containing or not containing fish, turtles, or crayfish added by the owner and the count of garden ponds with different frequency of occurrence of odonates, adult amphibians, and tadpoles as observed by the pond owner (0-never; 1-once; 2-sometimes; 3-often)

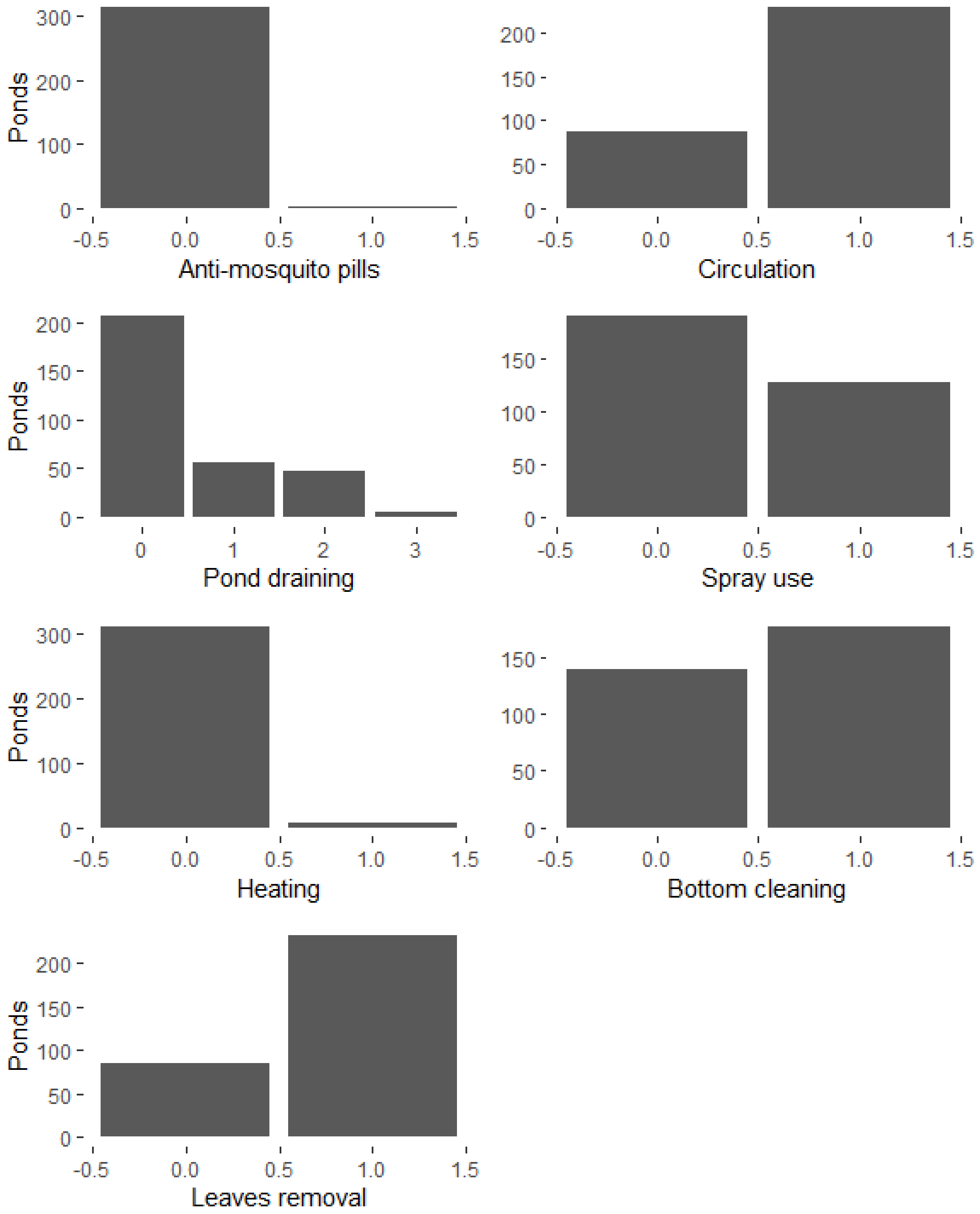


**Fig. S2** Bar plots showing the count of garden ponds with different frequencies of local management activities practised (0-never, 1-less than once a year, 2-once a year, 3-more than once a year). Anti-mosquito pills: the use of chemicals against mosquito larvae (pills); Circulation: presence of water circulation; pond draining; Spray use: the use of insecticide/pesticide spray in the garden; heating; bottom cleaning; leaves removal

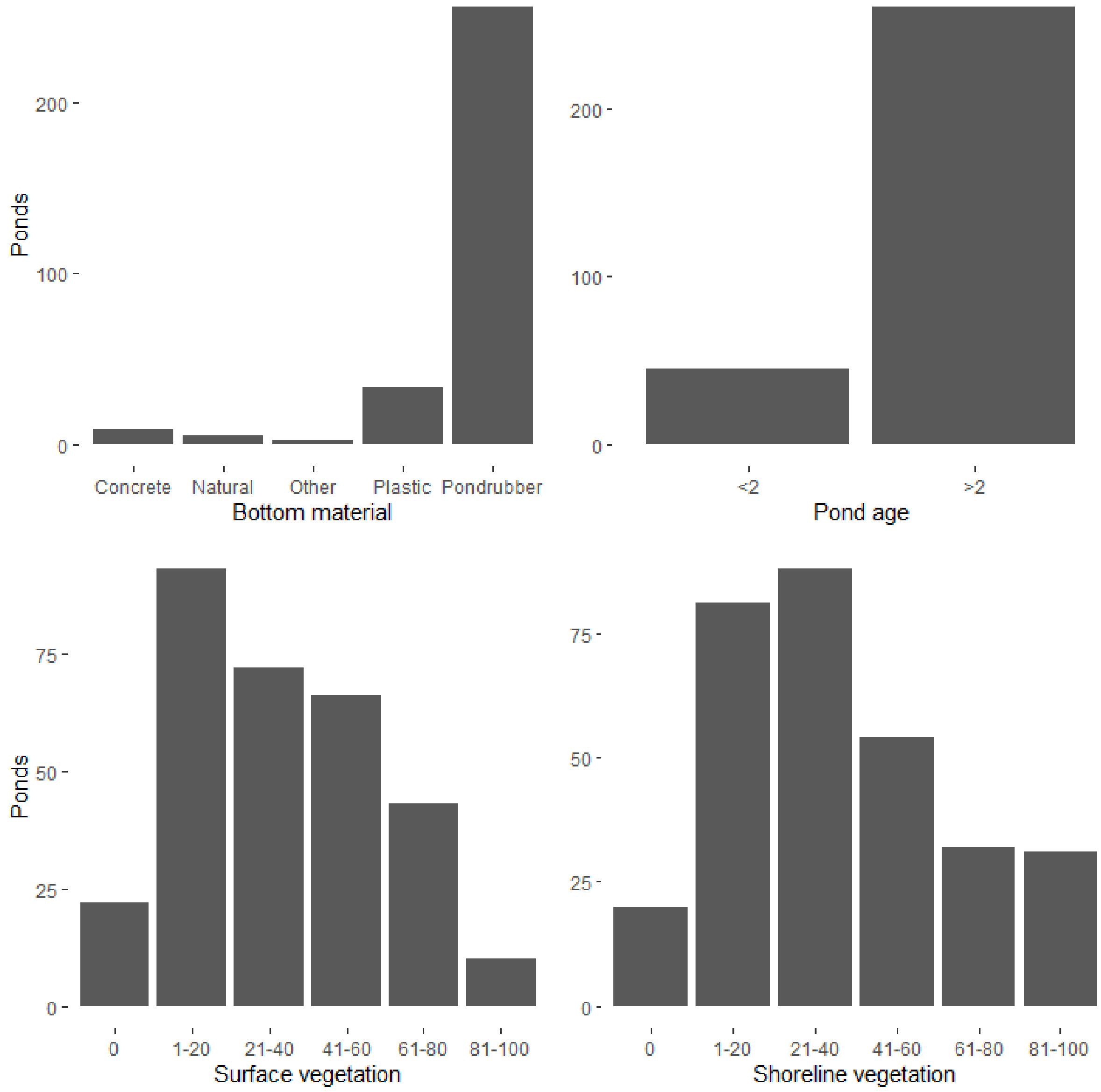


**Fig. S3** Bar plots showing the count of garden ponds within different categories of pond bottom material (concrete, natural, plastic, rubber pond liners, other), pond age (old ponds >2 years old and newly created ponds ≤2 years old), and surface and shoreline vegetation cover in % (0; 1-20; 21-40; 41-60; 61-80; 81-100)

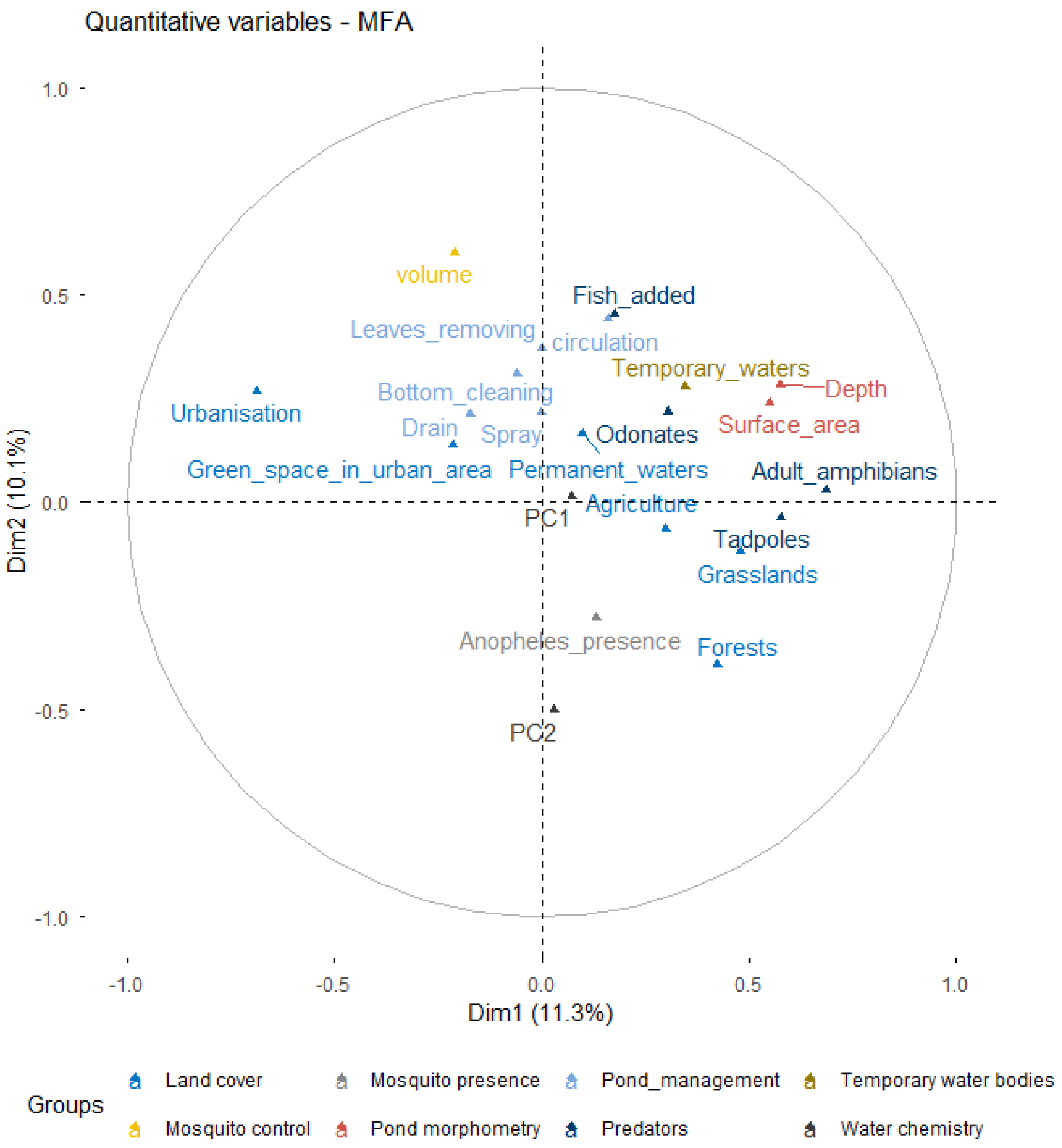


**Fig. S4** Multiple factor analysis (MFA) biplot showing the associations among land cover categories, environmental variables, and pond management activities, and the presence of *Anopheles maculipennis* larvae in garden ponds. Spray: use of insecticide/pesticide spray in the garden; drain: draining of garden ponds; volume: volume of chemical control of mosquitoes; PC1: first principal component (high negative loadings for heavy metals: Ni, Fe, Mn), PC2: second principal component (high negative loadings for pH and conductivity)

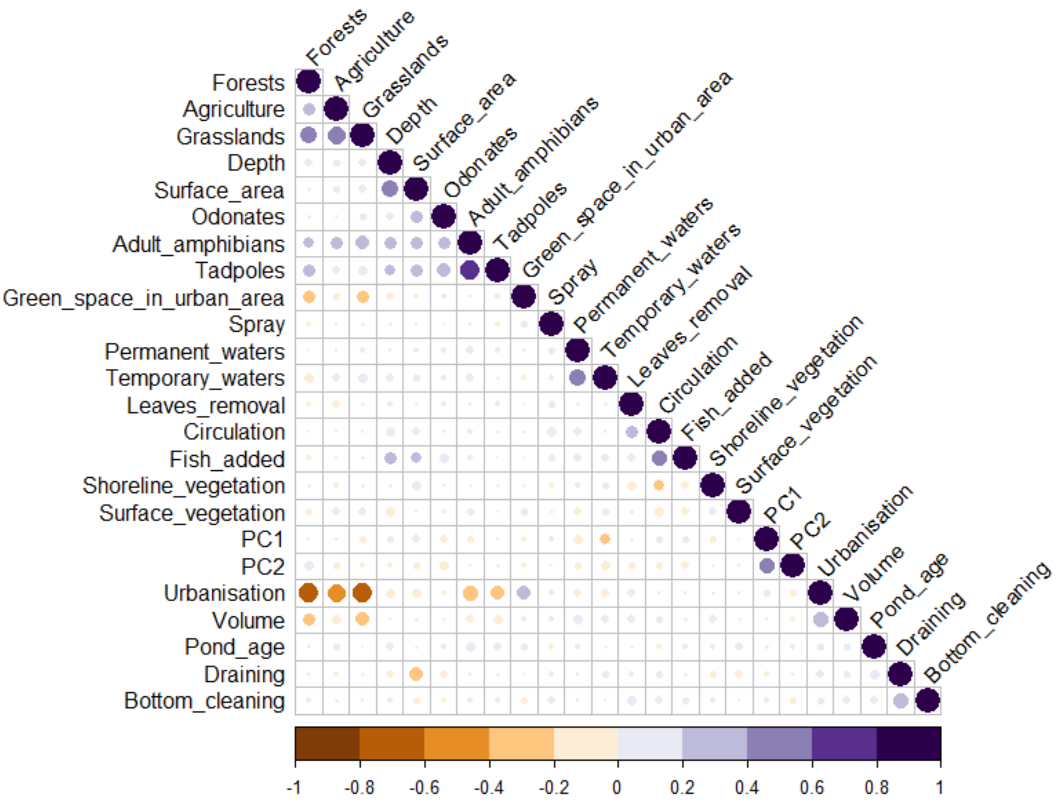


**Fig. S5** Correlation plot showing Spearman's rank correlations between each pair of predictor variables: urbanisation, green space in urban area, agriculture, grasslands, forests, permanent waters, depth, surface area, PC1 (high loadings for heavy metals: Ni, Fe, Mn), PC2 (high loadings for pH and conductivity), temporary waters, chemical mosquito control volume (Volume), pond draining, presence of water circulation, use of insecticide/pesticide spray in the garden (Spray), bottom cleaning, leaves removal, odonates, adult amphibians, tadpoles, fish, surface and shoreline vegetation, and pond age

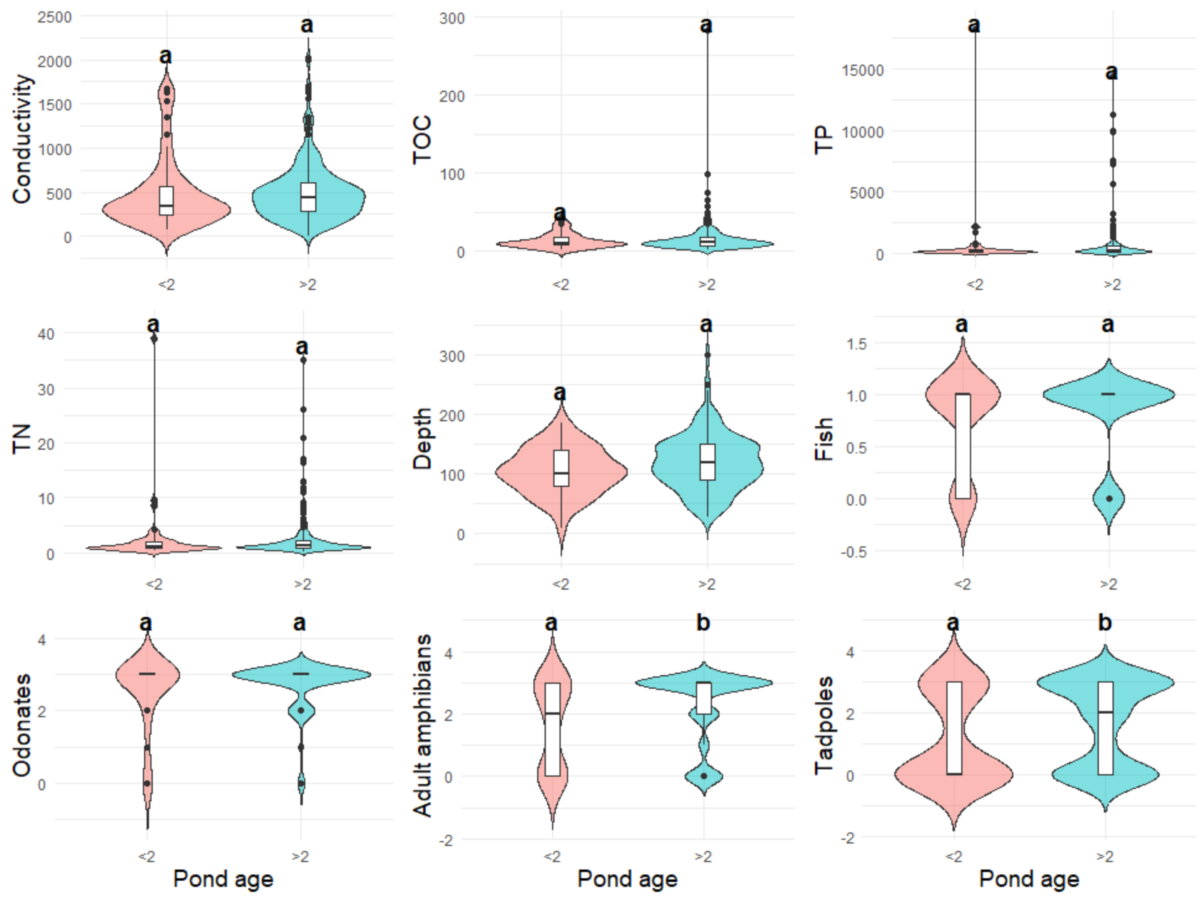


**Fig. S6** Violin plots showing differences of conductivity (µS/cm), total organic carbon (mg/L; TOC), total phosphorus (µg/L); TP), total nitrogen (mg/L; TN), pond depth (cm), presence/absence of fish (1/0), the frequency of occurrence of odonates, adult amphibians and tadpoles (0-never; 1-once; 2-sometimes; 3-often) between newly created garden ponds (≤ 2 years old) and older garden ponds (> 2 years old). Kruskal-Wallis test, followed by post-hoc Dunn's test was conducted, as the data did not meet the assumptions of normality. Significant differences between pond age categories ($P < 0.05$) are indicated by differing compact letter displays on the plot

**Table S1** List of generalised linear models testing the effects of selected predictors on the probability of the occurrence of *Anopheles maculipennis* larvae in garden ponds with their Akaike's Information Criterion (AIC) and Tjur's $R^2$ (coefficients of determination). The final selected model with the lowest AIC is highlighted in bold.

| Model | AIC | $R^2$ |
|---|---|---|
| Null model (no predictors) | 265.2 | 0 |
| Full model (Model 1)<br>(land cover + pond management + mosquito control + predators + water chemistry + pond age + vegetation + bottom material) | 267.9 | 0.154 |
| Model 2<br>(land cover + pond management + mosquito control + predators + water chemistry + pond age + vegetation) | 262.2 | 0.150 |
| **Model 3**<br>**(land cover + pond management + mosquito control + predators + water chemistry + pond age)**<br>***Model 3 without quadratic terms** | **257.5**<br><br>**252.6** | 0.102<br><br>0.099 |
| Model 4<br>(land cover + pond management + mosquito control + predators + water chemistry) | 264.4 | 0.069 |
| Model 5<br>(land cover + pond management + mosquito control + predators) | 268.3 | 0.065 |
| Model 6<br>(land cover + pond management + mosquito control) | 266.6 | 0.063 |
| Model 7<br>(land cover + pond management) | 267.2 | 0.055 |
| Model 8<br>(land cover) | 267.6 | 0.028 |

**Table S2** Results of the final selected generalised linear model for *Anopheles maculipennis* (AIC = 252.6; $R^2 = 0.099$). P values ≤ 0.05 are highlighted in bold. PC2: second principal component (high negative loadings for pH and conductivity).

| Predictors | Estimate | SE | Z value | P value |
|---|---|---|---|---|
| Intercept | -0.95 | 0.65 | -1.11 | 0.27 |
| Urbanisation | -0.01 | 0.01 | -0.80 | 0.43 |
| Green space in urban area | 0.01 | 0.01 | 1.07 | 0.29 |
| Agriculture | 0.06 | 0.02 | 2.51 | **0.01** |
| Bottom cleaning | -0.49 | 0.39 | -1.28 | 0.20 |
| Draining | 0.24 | 0.23 | 1.04 | 0.30 |
| Spray use | 0.63 | 0.35 | 1.78 | 0.07 |
| Leaves removing | -0.64 | 0.37 | -1.73 | 0.08 |
| Mosquito control volume | -0.00 | 0.00 | -1.41 | 0.16 |
| Tadpoles | 0.11 | 0.14 | 0.79 | 0.43 |
| PC2 | -0.01 | 0.14 | -0.10 | 0.92 |
| Pond age | -1.25 | 0.42 | -2.97 | **0.003** |
